# Glucose decoration on wall-teichoic acid is required for phage adsorption and InlB-mediated virulence in *Listeria ivanovii*

**DOI:** 10.1101/2021.03.08.434512

**Authors:** Eric. T. Sumrall, Stephan R. Schneider, Samy Boulos, Martin J. Loessner, Yang Shen

## Abstract

*Listeria ivanovii* (*Liv*) is an intracellular Gram-positive pathogen that primarily infects ruminants, but also occasionally causes enteric infections in humans. Albeit rare, this bacterium possesses the capacity to cross the intestinal epithelium of humans, similar to its more frequently pathogenic cousin, *Listeria monocytogenes* (*Lmo*). Recent studies in *Lmo* have shown that specific glycosyl modifications on the cell wall-associated glycopolymers (termed wall-teichoic acid, or WTA) of *Lmo* are responsible for bacteriophage adsorption and retention of the major virulence factor, Internalin B (InlB). However, the relationship between InlB and WTA in *Liv* remains unclear. Here, we report the identification of the unique gene, *liv1070* that encodes a putative glucosyltransferase in the polycistronic WTA gene cluster of the Liv WSLC 3009 genome. We found that in-frame deletion of *liv1070* led to loss of the glucose substitution on WTA, as revealed by UPLC-MS analysis. Interestingly, the glucose-deficient mutant became resistant to phage B025 infection due to an inability of the phage to adsorb to the bacterial surface, a binding process mediated by the receptor-binding protein B025_Gp17. As expected, deletion of *liv1070* led to loss of InlB retention to the bacterial cell wall, which corresponded to a drastic decrease in cellular invasion. Genetic complementation of *liv1070* restored the characteristic phenotypes, including glucose decoration, phage adsorption, and cellular invasion. Taken together, our data demonstrate that an interplay between phage, bacteria, and host cells also exists in *Listeria ivanovii*, suggesting the trade-off between phage resistance and virulence attenuation may be a general feature in the *Listeria* genus.

**Importance:** *Listeria ivanovii* is a Gram-positive bacterial pathogen known to cause enteric infection in rodents and ruminants, and occasionally in immunocompromised humans. Recent investigations revealed that, in its better-known cousin *Listeria monocytogenes*, strains develop resistance to bacteriophage attack due to loss of glycosylated surface receptors, which subsequently resulting in disconnection of one of the bacterium’s major virulence factors, InlB. However, the situation in *L. ivanovii* remains unclear. Here, we show that *L. ivanovii* acquires phage resistance following deletion of a unique glycosyltransferase. This deletion also leads to dysfunction of InlB, making the resulting strain unable to invade host cells. Overall, this study suggests that the interplay between phage, bacteria and the host may be a feature common to the *Listeria* genus.

## Introduction

The *Listeria* genus contains two species that are capable of causing disease in mammals. *L. monocytogenes* (*Lmo*) is by far the more common of the two, which is capable of invading and replicating within mammalian cells, and has a mortality rate of up to 30% (1). While *Lmo* is capable of causing disease in both animals and humans, *L. ivanovii* (*Liv*) does so almost exclusively in ruminants (2) and rodents (3). *Liv* has on occasion demonstrated its ability to cause disease in humans (4–7), showing that while it may be less adapted to cause established infection in humans, it possesses all the necessary virulence factors and has the capability given the right patient circumstances. There are two subspecies within *Liv*, including spp. *ivanovii* and spp. *londoniensis*. Both have been described and categorized based on the ability to metabolize *N*-acetyl-mannosamine and ribose. *Liv* spp. *ivanovii* is generally sensitive to infection by many phages, while strains of spp. *londoniensis* appear to be quite resistant to phage attack due to the presence of a functional type II-A CRISPR-Cas system (8). To date, only *Liv* spp. *ivanovii* has been shown to cause listeriosis in human and animals (3, 9). Like *Lmo*, *Liv* infections are thought to be food-borne, as bacteria have been isolated from both the feces and blood from infected human patients (4). From this, it can be hypothesized that *Liv* also possesses the capability to cross membrane barriers during infection, and indeed, has been shown in cell culture to be internalized in both human and bovine cell lines, and possesses the major pathogenicity island similar to that in *Lmo* (6). *Liv* is also thought to be rarer in the environment, suggesting that host tropism and lower pathogenicity may not be the only reasons for its infrequent disease occurrence.

All species in the *Listeria* genus can be characterized by serotyping, a serological process that relies upon the structural variation of the bacterial cell surface. Of the many serovars (SVs) within the *Lmo* species, 1/2 and 4b cause the vast majority of disease in humans, suggesting that there may be a clinical relevance for the surface structures that confer serovar identity (10), which are primarily the wall-teichoic acids (WTAs) (11). WTAs are complex carbohydrate molecules that make up a majority (up to ~60%) of the dry weight of the bacterial cell wall and are covalently conjugated to the peptidoglycan and extend outward (12). Beyond serving as the major antigenic determinants for serotyping, WTAs are involved in several key physiological functions, including maintaining osmotic pressure, antibiotic resistance, virulence, and interaction with host cells and bacteriophages (13–18). WTAs consist of a single glycosylated glycerol-based linkage unit, and a chain of 20-30 repeating units which can vary in structure between individual strains (19). In *Listeria*, the WTA repeating units are generally made up of ribitol phosphate (type I WTA), but in some serovars contain an *N*-acetylglucosamine (GlcNAc) residue integrated into the chain (type II WTA) (20). Further structural variation stems from the carbon position at which GlcNAc is linked to the ribitol residue, or further glycosylation or other modifications, termed “decorations” (20). *Liv* strains are designated SV 5, and feature an integrated GlcNAc residue conjugated to the C2 position of the ribitol in the primary WTA chain. This GlcNAc is further decorated with a glucose residue in most cases, but the chain can contain interspersed repeating units that are *O*-acetylated, or not modified at all (21).

Work in our lab with *Lmo* showed that in the highly virulent SV 4b strains, bacteriophage predation can select for resistant strains that lack a galactose (Gal) decoration on their WTA, as certain bacteriophages specifically recognize Gal for their adsorption (22). This consequently led to an inability of the mutant strains to invade host cells and a drastic reduction in virulence, due to the fact that the major virulence factor, Internalin B (InlB), relies upon this Gal modification for its retention to the bacterial surface (22). InlB is well-established as playing a large role in the invasion of the host liver, spleen and placenta by recognizing the cMet receptor and inducing the endocytic pathway (23, 24). We showed that the InlB virulence factor relies exclusively on WTA, and not lipoteichoic acid (a similar, membrane-anchored glycerol-based polymer), which was previously thought to be responsible for InlB’s surface retention (25). Similar studies in *Lmo* SV 1/2 showed that InlB relies on the rhamnose residue on WTA for its surface retention (26), a decoration that can also function as a phage receptor (27). Because these decorations appear to serve both as receptors for bacteriophage binding, as well as ligands for the surface-associated virulence factor InlB, it became evident that *Listeria* faces a tradeoff between maintaining virulence and developing bacteriophage resistance (18). Because *Liv* genomes also feature *inlB* (6), we set off to determine whether the glucose (Glc) decoration on the WTA of this unique species plays a role in InlB retention and function, and whether like *Lmo*, *Liv* must also face a tradeoff between being resistant to bacteriophage predation, or maintaining a primary virulence function.

## Results

### Organization of the WTA biosynthesis gene cluster in ***Liv*** WSLC 3009

The genome of WSLC 3009 has been recently sequenced. Since much is known about the genetic characteristics and function of WTA in rod-shaped bacteria, we identified and annotated the WTA gene cluster in 3009 using the prototype polyribitol phosphate WTA-producing strain *B. subtilis* W23 (28). The majority of polycistronic WTA genes are in a single locus (Fig. **1A**), in addition to the monocistronic tarO and tarA genes, whose transcribed products are known to initiate WTA biosynthesis (29). In this locus, the function of most genes can be predicted based on sequence similarity to their homologs in *B. subtilis*, *S. aureus*, and *L. monocytogenes*. The sequential action of TarO, TarA, and TarB enzymes produce the conserved GlcNAc-ManNAc-Gro linkage unit (28). TarI, TarJ, and TarD produce and transfer the CDP-ribitol substrate involved in the WTA glycosylation process (30). Liv1073 possesses a GT-2 (type 2 glycosyltransferase) domain, and it is conserved across *Listeria* subspecies (Fig. **S1**) featuring type II WTA with an integrated GlcNAc in the primary WTA chain (20). In our hands, previous attempts to delete the Liv1073’s homolog in 4b *Lmo* did not yield a viable mutant, which led us to speculate that this gene is involved in the addition of GlcNAc onto the polymer chain, a process required for cell growth and development. Other members of this gene cluster are annotated OatT (22) (47% AA sequence identity), IspC (31) (limited sequence identity, but contains conserved glycine-tryptophan modules and the amidase motif), GttA (25) (34% AA identity), and GalU (89% AA identity), which have recently been described in *Lmo* SV 4b (22). Domain homology searches did not reveal any functional domains in Liv1067, yet Liv1070 was found to be conserved in *L. ivanovii* and some other *Listeria* subspecies (**Table. S1**). The encoded protein bears an *N*-terminal GT-2 domain (pfam00535), and a glycerol phosphotransferase domain at its *C*-terminus (Fig. **1B**). We therefore hypothesized that Liv1070 is responsible for the glucose decoration on WTA (Fig. **1C**). To test this hypothesis, we first generated the knock-out mutant 3009Δ*liv1070* and found that this mutant strain did not show any growth defects relative to the WT strain (Fig. **1D**).

**Figure 1.**
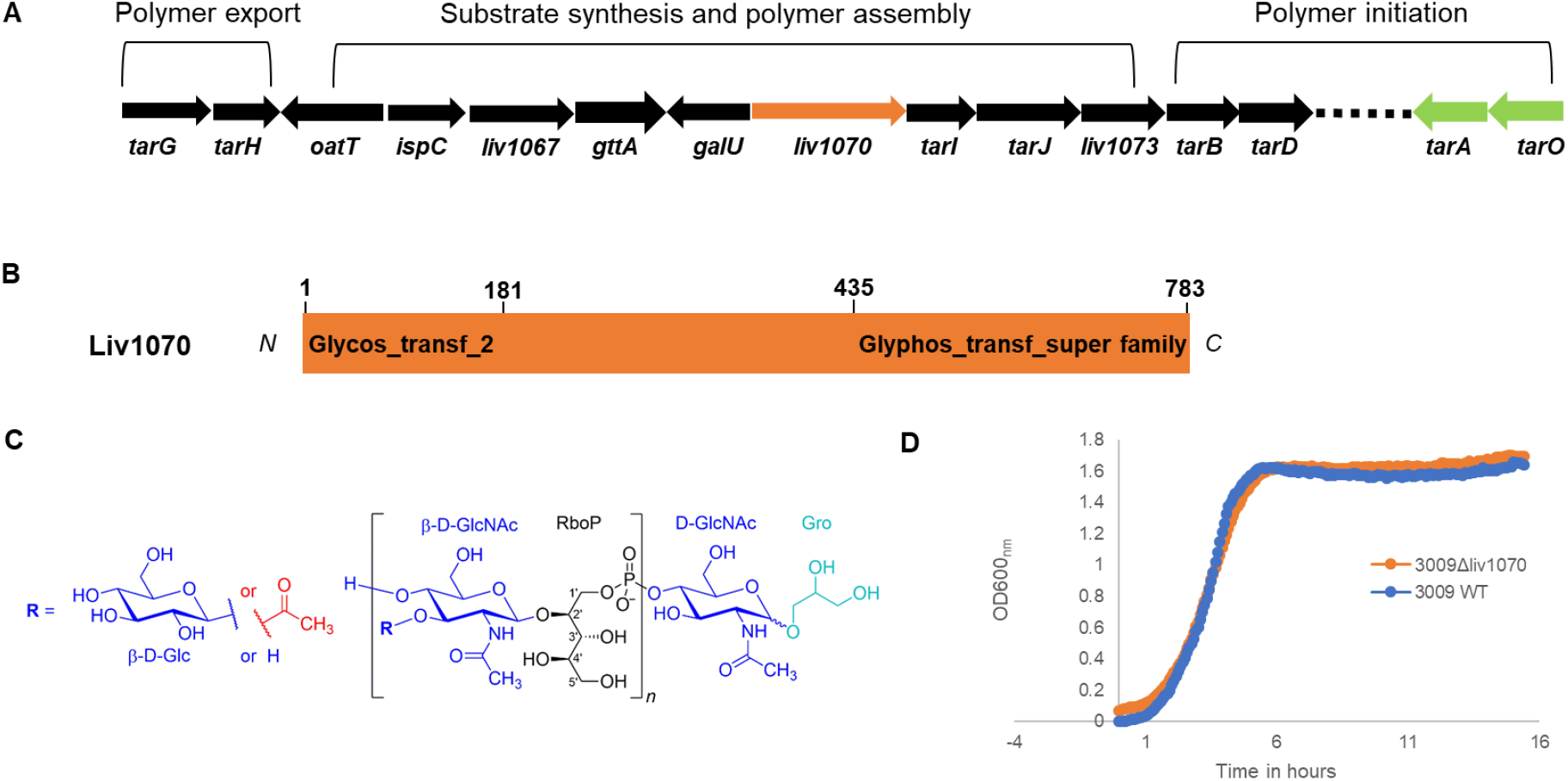
Genetic organization of *L. ivanovii* WSLC 3009 wall teichoic acid biosynthetic genes. (A) The ~18.8 kb WTA locus is denoted in black arrows. Dash line indicates the gap. The annotated function of respective genes is shown above the locus. (B) Domain architecture of Liv1070 based on conserved domain search. (C) Chemical structure of 3009 WTA repeating and linkage unit. Abbreviations: Glc, glucose; GlcNAc, *N*-acetylglucosamine; Rbo, ribitol; Gro, glycerol; P, phosphate. (D) Growth curves of indicated strains, as determined by measuring the OD600_nm_ over the course of 16 hours. The results from one representative are shown.

### In-frame deletion of *liv1070* results in loss of glucose decoration on WTA

To further verify that *liv1070* confers WTA glucosylation in the parent strain 3009, WTA was purified from the WT and mutant strains using previously-described analytical technique (20). The structure of the WTA repeating unit was determined using ultra-performance liquid chromatography, coupled to mass spectrometry (UPLC-MS) (Fig. **2**). The structure of the WT strain showed two major peaks, one with m/z 354 (representing the GlcNAc-Rbo fragment), and the other at 516 (with the Glc decoration) (see Fig. **2**). As is consistent with previous findings, this shows in the WT 3009 strain that only a portion of WTA repeating unit structures are glycosylated, in this case with glucose. In the 3009Δ*liv1070* strain however, the peak with m/z 516 is completely missing, indicating that no Glc decoration exists on the WTA of this strain. As expected, the chromatogram of the 3009Δ*liv1070*∷pPL2(*liv1070*) complemented strain (see Methods section) appears identical to the WT strain, indicating that the phenotype is fully restored.

**Figure 2.**
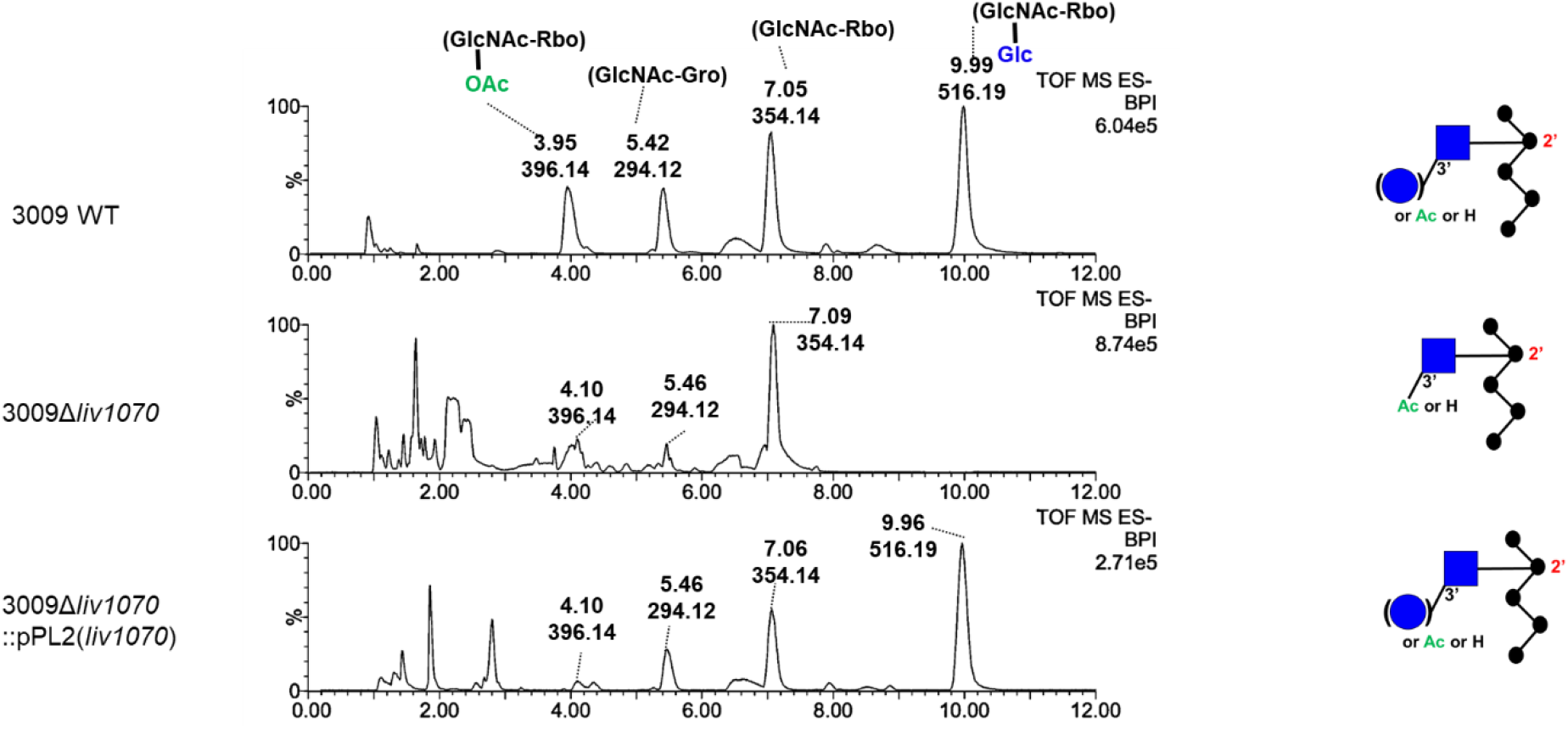
Liquid chromatographic separation and mass-spectrometry identification of WTA repeating unit of the three indicated strains, as determined by UPLC-MS. Major peaks are labeled with their respective retention time (Rt; [min] and base peak ion [M-H]^−^ (*m/z*)). The deduced structures of the respective WTA type with C2 *N*-acetyl substituent are shown on the right in CFG representation (Consortium for Functional Glycomics). Abbreviations: Glc, glucose (blue circle); GlcNAc, *N*-acetylglucosamine (blue square); Rbo, ribitol; Gro, glycerol; P, phosphate; OAc, *O*-acetyl group.

### Loss of glucosylated WTA renders *Liv* 3009 cells insensitive to phage adsorption

Bacteriophage B025 is thought to utilize the Glc decoration on the WTA for binding and recognition, as it features a specificity for SV 5 *Liv* strains. To determine whether deletion of *liv1070* confers phage resistance, a pulldown assay was performed. As expected, far fewer B025 phage particles adsorbed to the surface of the 3009Δ*liv1070* strain (Fig. **3A**). To further verify that the Glc moiety was missing from the WTA in strain 3009Δ*liv1070*, the fluorescently labeled receptor binding protein B025_Gp18-GFP was utilized for glycotyping (32, 33). We previously demonstrated that this protein binds and recognizes *Listeria* strains possessing WTA monomers with the GlcNAc moiety linked to the C2 position of ribitol, and decorated with Glc or Gal (33). Thus, loss of Glc decoration from the WTA of strain 3009 would lead to an inability of the protein to recognize the bacterial cell surface. Indeed, incubation of 3009Δ*liv1070* with the B025_Gp18-GFP fusion protein showed no fluorescence signal relative to that of the 3009 WT strain (Fig. **3B**). The complemented strain showed a restored B025_gp18-GFP protein binding and phage adsorption, demonstrating that *lmo1070* is sufficient to confer this phenotype. Together, these data strongly suggest that the gene *lmo1070* is responsible for WTA glucosylation in the *Liv* strain 3009, a structure which mediates bacteriophage adsorption and susceptibility.

**Figure 3.**
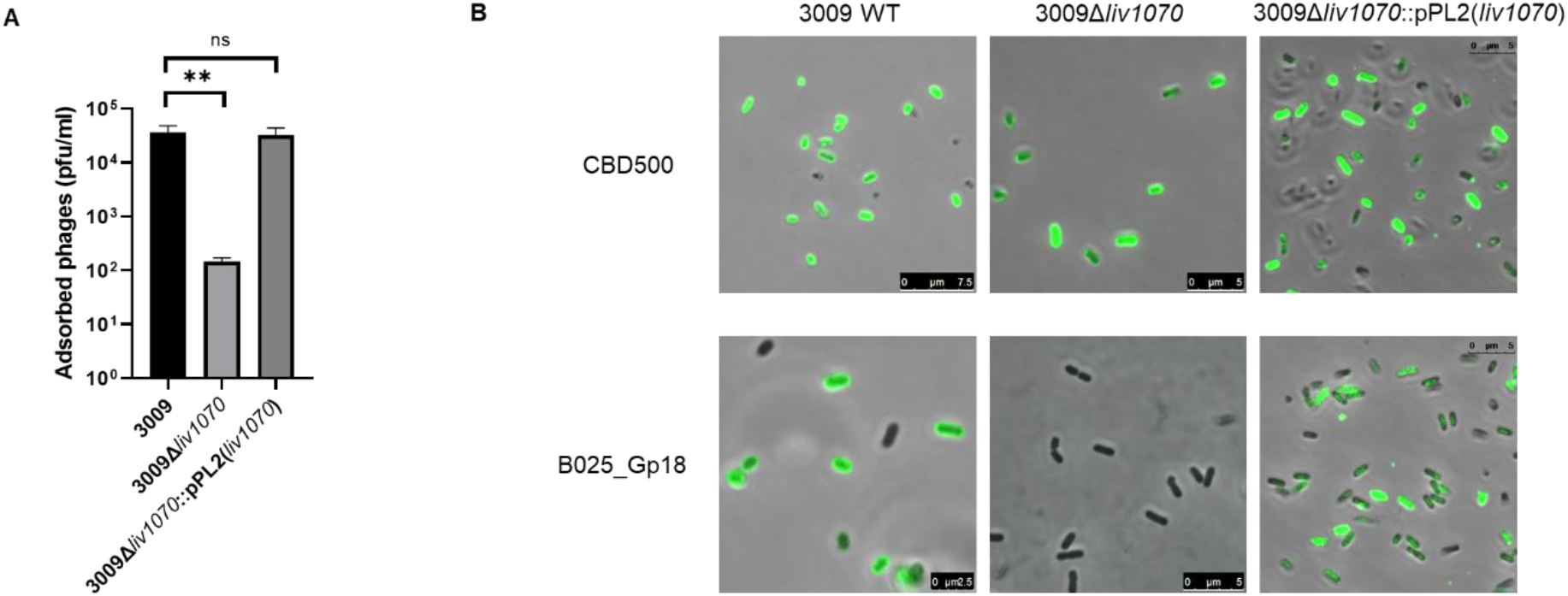
WTA glucosylataion in *L. ivanovii* determines phage adsorption via its interaction with the receptor-binding protein. (A) Total adsorbed Listeria B025 phages, as determined by a pulldown assay and quantification by PFU counting (bars represent mean +/− SEM). b) Staining of the indicated Liv strains with the phage-derived recombinant CBD-binding protein CDB500, and the receptor-binding protein B025_Gp18, both fused to GFP. Visualization performed using fluorescence confocal microscopy to detect GFP signal. For both assays, experiments were performed in triplicate. **P <0.01

### Glucosylated WTA is required for InlB cell wall association and Caco-2 and Hela cell invasion

Because *Liv* is an invasive species causing disease in certain animals, their cells likely harbor functional InlB on the surface. InlB has been shown to rely upon WTA rhamnose decorations for its surface retention in SV. 1/2 strain in *Lmo* (26). In SV 4b strains, it is also known that InlB relies upon the WTA galactose decoration for its surface retention, but not the glucose decoration (25). *Liv* possesses a similar WTA structure to that of the *Lmo* SV 4b, but differs in its GlcNAc connectivity, and features only Glc decoration instead of both Glc and Gal in SV 4b. We thus hypothesized that Glc alone may be responsible for InlB retention via *Liv* type II WTA.

To evaluate whether the loss of the Glc decoration on the WTA of strain 3009 affects InlB surface retention, Western blot assays were performed using whole-cell protein extracts and precipitated supernatant, and tested with anti-InlB antibody. As can be seen, the 3009Δ*liv1070* strain seems to lose the surface-associated InlB protein (Fig. **4A**), similar to what has been described for the *gttA* mutant in SV 4b *Lmo (22)*. The phenotype was restored in the complemented 3009Δ*liv1070*::pPL2(*liv1070*) strain. Overall, it appears that 3009 expresses lower levels of InlB protein than *Lmo* strain 1042 (Fig. **4A**), which may be consistent with *Liv* being a somewhat less virulent species. To determine whether the loss of InlB in this Glc-deficient mutant has an effect on the strain’s ability to invade host cells, a gentamicin protection assay was performed in both HeLa cells and the Caco-2 epithelial cell line (Fig. **4B**). Because the 3009Δ*liv1070* strain is severely deficient in its invasive abilities, it can be assumed that InlB function is lost. This is supported by the observation that invasion into HeLa cells was almost abolished. As HeLa cells do not express E-cadherin, invasion in this cell line is known to be entirely InlB-dependent (34). The low invasion levels of 3009Δ*liv1070* in Caco-2 cells, which do express E-cadherin, is presumably mediated via the InlA pathway. Invasion of the complemented strain 3009Δ*liv1070*::pPL2(*liv1070*) is insignificantly different from the WT strain, demonstrating that the Glc decoration on WTA is sufficient to maintain proper invasion levels via its ability to retain InlB on the cell surface. Together, these data clearly show that glucosylation of the WTA in SV 5 *L. ivanovii* mediates phage resistance and maintains the function of one of the major *Listeria* virulence factors.

**Figure 4.**
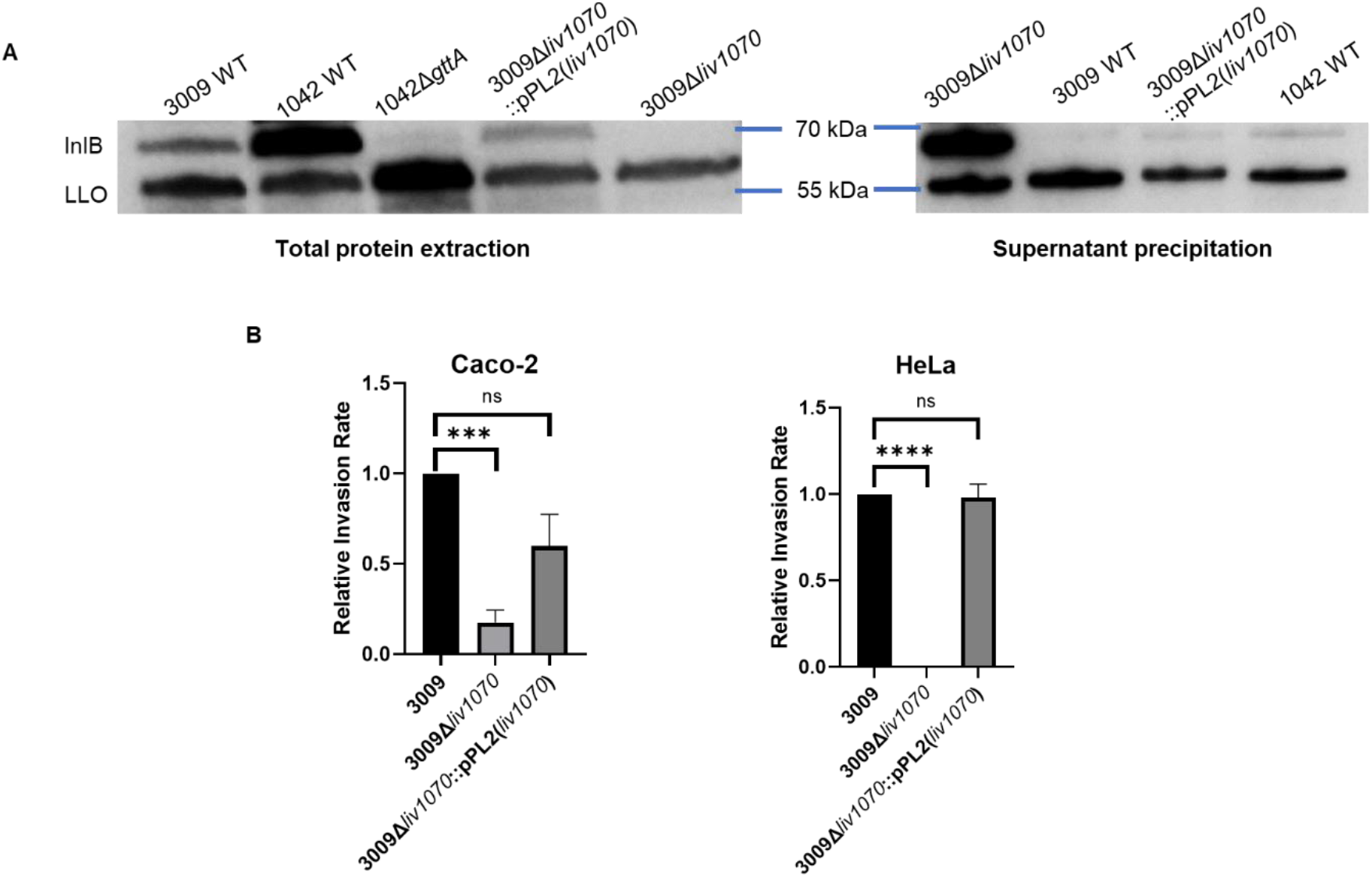
*L. ivanovii* requires glucosylated teichoic acids for InlB-mediated invasion. (A) Western blot of *Liv* total and surface proteins of indicated strains, detected using an anti-InlB antibody with anti-LLO (Listeriolysin) as a loading control. *L. monocytogenes* serovar 4b strain WSLC 1042 and its Δ*gttA* KO derivative were compared to demonstrate the relative or decreased amount of surface-associated InlB. (B) Relative invasiveness of the indicated strains compared to that of *Liv* 3009 WT, in both Caco-2 and HeLa cell lines for 3 hours (mean normalized to 3009 WT ±SEM, as determined by a gentamicin protection assay. For both assays, n=4.***P<0.001; ****P<0.0001; ns, not significant relative to 3009).

## Discussion

In this investigation, we sought to determine whether the InlB virulence factor also relies upon WTA decoration, and more specifically glucosylation in *L. ivanovii*. Previous investigations in *L. monocytogenes* showed that InlB requires presence of galactose on the WTA polymer for its surface retention, and this decoration is also essential for phage recognition and binding (22, 25, 26). However, *L. ivanovii* possesses a somewhat different WTA structure, with a different type of glycosylation, a structure which confers the unique SV 5 designation (20). Here, we show that the gene *liv1070* is necessary for glucosylation of the WTA monomer. Its deletion led to the loss of glucosylation, which was determined both structurally and by glycotyping with specific WTA-binding phage proteins (33). As expected, the glucose moiety is utilized by phage B025, which specifically infects strains possessing glycosylated WTA with an integrated GlcNAc linked to the ribitol backbone at the C2 position. Loss of glucosylation confers phage resistance and leads to a loss of surface InlB, together showing that like *Lmo*, *Liv* must also face an evolutionary tradeoff: to maintain an important virulence factor that mediates invasion of certain cell types, or be resistant to predation by bacteriophages.

The WTA of *Lmo* SV 4b cells possesses both a glucose and galactose decoration, but it was shown that only the galactose and not glucose decoration was responsible for InlB surface retention (25). The data shown here strongly suggest that the conserved InlB SH3b domain, which is the domain responsible for WTA binding, evolved to specifically recognize different glycosylated WTA types (22, 26). It may be the position and orientation of the sugar moiety on the WTA that governs binding to InlB, as it has been shown that glucosylated GlcNAc at the C4 position of ribitol or galactosylated GlcNAc linked at the C2 position failed to retain InlB on the bacterial surface (22). How InlB interacts with different types of WTA is still not fully understood, but it will be interesting to experimentally test in future specificity studies. Data have suggested that most of the protein is buried within the bacterial cell wall, and only a small portion is available to the outside. This can be demonstrated when preparing *Lmo* for immunofluorescence, as the cell wall has to be partially digested with lysozyme in order for the anti-InlB antibody to access its epitope (22). Evidence suggests that the SH3b domain of InlB, which sits at the *C*-terminus at the protein (the *N*-terminus contains the membrane-spanning portion of the protein), and may explain why it interacts with the WTA, but not the LTA, which is likely more buried within the cell wall (25, 35). However, activation of the host cMet receptor requires both the SH3b domain as well as the LRRs, which are more *N*-terminal, and thus further buried within the cell wall (36). How it is that InlB, a membrane-associated protein which may only be partially exposed above the surface of the cell wall activates the host cell receptor, is still not fully understood and requires further study.

InlB functions by recognizing the host cell receptor cMet, and inducing the endocytic pathway. Data presented here have shown that the phage-resistant strain lacking a WTA glycosyltransferase does not express InlB on its cell surface and is consequently deficient in its ability to invade host cells. InlB is involved in the invasion of the liver, spleen and placenta (23, 37–39), meaning that its loss leads to a virulence attenuation. *Lmo* strains deficient in WTA glycosylation lack the function of other proteins, namely ActA (40), and the autolysins (26). Whether this is also the case in *Liv* remains to be seen, but it can be theorized that the glucose-deficient 3009Δ*liv1070* discussed here would show a significant virulence attenuation in an animal model of *Liv* infection, as it has been shown that *Liv* can heavily colonize in the liver (41).

We have identified a putative enzyme (Liv1070) required for the glucosylation of WTA in *Liv*. Based on *in-silico* predictions and the previous findings obtained in *Lmo* (25), we propose a model (Fig. **5**) for the WTA glucosylation process in *Liv*. In previous work, GttA was shown to be involved in catalyzing the addition of Gal onto the undecaprenyl lipid carrier (C_55_-P). Due to the close homology to GttA, Liv1068 is thus predicted to be a cytoplasmic glycosyltransferase that transfers UDP-Glc to the lipid carrier. In addition, we presume that Liv2596 functions as a putative flippase (84% identity to GtcA) that transports the C_55_-P-Glc lipid intermediate across the membrane (42). We show that 3009Δ*liv1070* lacks Glc decorations on WTA, suggesting that *liv1070* might encode a GT-B fold glucosyltransferase responsible for the transfer of the Glc residues onto the WTA chain. Since Liv1070 is predicted to feature 2 transmembrane helices (43) (AA 121 to 141 and AA 585 to 713), we speculate that this glucosylation process may also occur on the outside of the cell as revealed in *Lmo* (44). Nevertheless, further biochemical evidence is still required to elucidate this process.

**Figure 5.**
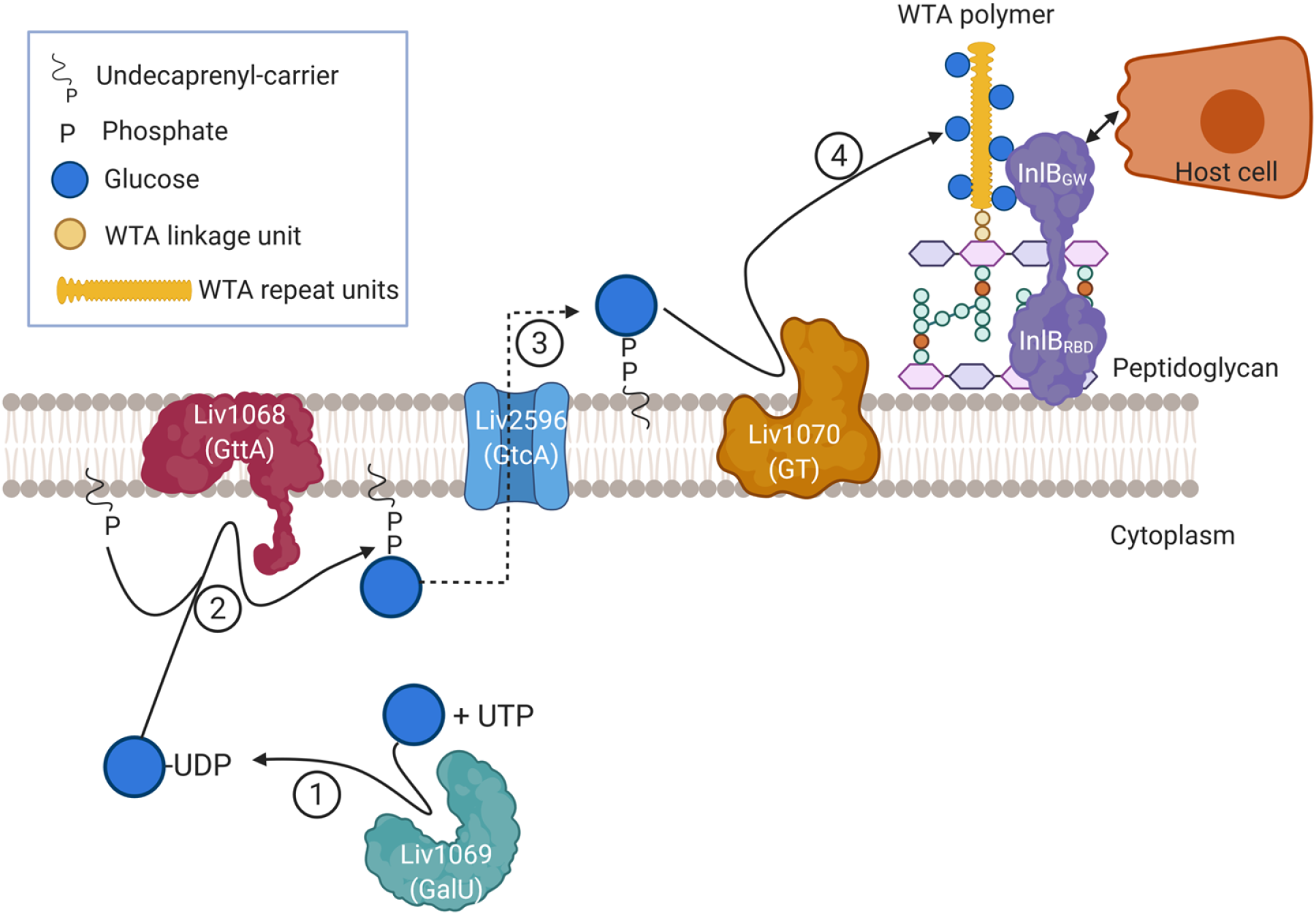
Proposed model for the WTA glucosylation process in *L. ivanovii*. Based on the bioinformatic and genetic data presented in this study, we propose that WTA in *L. ivanovii* is glucosylated with the aid of cytoplasmic GT Liv1069 (GalU, glucose-1-phosphate uridyltransferase), which we predict produces UDP-Glc from UTP and glucose-1-phosphate (step 1). Next, we hypothesize that Liv1068 catalyzes the addition of UDP-Glc residues onto the undecaprenyl lipid carrier (C_55_-P) on the inner leaflet of the cell membrane due to its high homology to GttA (step 2) as previously described in serovar 4b *L. monocytogenes*. The C-55-P-glucose intermediate is then transported across the membrane by Liv1069 (step 3), which is 84% identical to the *L. monocytogenes* GtcA protein reported in a previous study. The glucose residues are subsequently transferred onto the growing WTA chain outside the cell (step 4), and we suggest that this step is catalyzed by glucosyl transferase Liv1070. The WTA chain is conjugated by TarTUV homologue to the MurNAc of peptidolycan following export by an ABC transporter TarGH. The glucosylated WTA confers the retention of the InlB on the *Listeria* surface, which interacts with the host receptor cMet to activate receptor-mediated endocytosis, facilitating entry into host cells. Figure S1. Comparison of genes encoding type II WTA biosynthesis pathways in the indicated *Listeria* serovars and strains.

*Liv* is thought to seldomly occur in the environment. In the dairy industry, a study from Ireland showed that *Liv* isolates exist at a low prevalence of 1.4%, but these strains were similarly capable (or even better) at invading Caco-2 cells compared to *Lmo* EGDe (45). While the prevalence is lower than what is typically found for *Lmo* (46), it seems significant nonetheless. This also suggests that *Liv* may be more prevalent in the environment and in food processing plants than previously assumed, promoting a need for further research into approaches for its containment. Bacteriophages have evolved as a viable option for biocontrol of *Listeria*, although as with classical antimicrobials, the occurrence of resistant strains may present a significant hurdle. However, evidence from this study and other recent studies has shown that resistance to bacteriophages may often be accompanied by physiological and virulence defects, which could present benefits to the host and decrease pathogenicity (47, 48). This could further the argument for bacteriophage use in biocontrol measures, if it holds true that resistant strains are less harmful.

## Methodology

### Bacterial strains, plasmids, phages and growth conditions

All bacterial strains, plasmids and phages used in this study are listed in **Table S2**. *E. coli* XL1-Blue (Stargene) used for cloning and plasmid construction was routinely cultured in Luria–Bertani (LB) broth at 37°C. The *E. coli* BL21-Gold strain was used for protein expression. L. monocytogenes strains were grown in 1/2 brain heart infusion (BHI), at 30°C with shaking when working with phages, or at 37°C when the cells were used for infection studies. For a list of strains, plasmids and primers utilized in the study, see **Table S2**. Propagation and purification of bacteriophage B025 were performed using *L. ivanovii* strain 3009 as previously described (25). For growth curve determination, overnight cultures were diluted in full BHI to an OD of 0.05 in triplicate in a 96-well plate, and the OD_600nm_ was measured for 16 h in a plate reader set to 37°C.

### Production of mutant and complemented strains

Deletion knockouts were produced via allelic exchange. 500bp flanking regions of the gene *liv1070* from strain 3009 were produced by PCR, along with the pHoss1 plasmid backbone. The pHoss1 plasmid contains a temperature-sensitive origin of replication, which cannot replicate at higher temperatures. The primers used for this are listed above in **Table S2**, and contained homologous overlap regions to allow for assembly. The three fragments were assembled together using Gibson assembly, followed by transformation into *E*. *coli* XL1-Blue. The pHoss1 plasmid was extracted from a single colony of XL-1 blue containing the two 500bp flanking regions of gene *liv1070*, and transformed into *Liv* strain 3009. This transformant was grown at permissive temperatures, allowing for gene deletion to proceed via allelic exchange. The complete deletion of *liv1070* was confirmed by PCR and Sanger sequencing. Complementation of 3009Δ*liv1070* with a functional copy of *liv1070* was performed by using the chromosome-integrating vector pPL2 (49). The *liv1070* gene was inserted by Gibson assembly into plasmid pPL2 under the control of the native promoter.

### Phage pulldown assay

Pull down assay using bacteriophage B025 was performed using MOI 0.01 as previously described (22), using WSLC 3009 as the propagation strain. Following serial dilution of the phage, the number of phages adsorbed to the bacterial surface was evaluated by phage overlays and expressed as the total plaque-forming units of phage adsorbed to WT, mutant and complemented strain.

### *Listeria* glycotyping assay

The abilities of GFP-tagged CBD500 and B025_Gp18 to bind to *Listeria* cell surface were tested using a fluorescence binding assay as previously reported (33). Briefly, *Listeria* cells from log phase cultures were harvested by centrifugation, and resuspended in 1/5 volume of PBS (pH 7.4). 100 μl of cells were incubated with 5 μl of 1 mg/ml of GFP-CBDs and incubated for 5 min at room temperature. The cells were spun down, washed three times, and finally resuspended in PBS buffer. The samples were then subjected to confocal laser scanning microscopy.

### WTA extractions and structural analysis

WTA was purified from the indicated *L*. *ivanovii* strains as previously described (20). Purified WTA polymers were depolymerized into monomeric repeating units by hydrolysis of the phosphodiester bonds using 48% hydrofluoric acid for 20 h at 0°C. The purified WTA monomers were lyophilized and subjected to UPLC-MS/MS for compositional and structural analysis.

### Gentamicin protection assay

The Caco-2 and HeLa cell lines used for *in vitro* assays were cultured at 37°C with 5% CO_2_ in DMEM GlutaMAX (Gibco), supplemented with sodium pyruvate, 1% non-essential amino acids and 10% FBS. Before infection, human cells were diluted to a concentration of 2-4×10^5^ cells/mL and seeded onto 96-well plates in triplicate the day before performing the experiment. On the day of infection, bacteria were grown at 37°C to an OD_600_ of 0.8–1.0, before being washed twice in PBS and diluted in DMEM lacking FBS to OD_600_ 0.01, which corresponds to a multiplicity of infection (MOI) of ~100. Human cells were washed twice with DPBS and bacterial suspensions were added on top. The co-culture was incubated for 2 h at 37°C. Cells were washed twice with DPBS and incubated a further 1 h in normal growth medium (with FBS) supplemented with 40 μg/mL gentamicin. Cells were lysed with 0.5% Triton-X-100, serially diluted, and 10 μL were spot-plated onto BHI agar plates. CFU counts were determined the following day. The number of bacteria that had adhered or invaded was expressed as a fraction relative to the invasion rate of strain WSLC 1042, which was normalized to 1 for each replicate.

### Western Blotting

To detect total InlB, total cell extracts were used instead of surface protein extracts in order to obtain a better visualization for the loading control. One mL of overnight culture grown in 1/2 BHI medium was mixed with 0.5 mL of 0.5 mm glass beads in a 2 mL tube and shaken on a vortex at maximum speed for 30 minutes. The tubes were spun down for 1 min at 200 x g and 500 μL of supernatant was transferred to a 1.5 mL Eppendorf, and spun again for 15 min at maximum speed. The supernatant was discarded and the pellet resuspended in 50 μL SDS sample buffer for an initial OD of 2 (volume was adjusted accordingly if initial OD varied) containing 5% β-mercaptoethanol, and boiled for 5 min. Samples were loaded onto an SDS-PAGE gel and western blot was performed as previously described (22), using a custom anti-InlB rabbit polyclonal antibody (1:5000), using a LLO antibody as the loading control (1:5000; Abcam. Inc). Supernatant extracts were produced via a TCA precipitation method using the supernatant from a 5mL culture, as previously described.

## Acknowledgements

This work was supported, in part, by Swiss National Science Foundation Grant 310030_156947/1. Graphic illustration was created using BioRender.com.

## Author Contributions

Conception and design: YS. Acquisition, analysis and interpretation of the data: ETS, SRS, SB, MJL, YS. Writing of the manuscript: ES, SRS. Reviewing and editing of the manuscript: MJL, YS.

---

**Figure S1.**
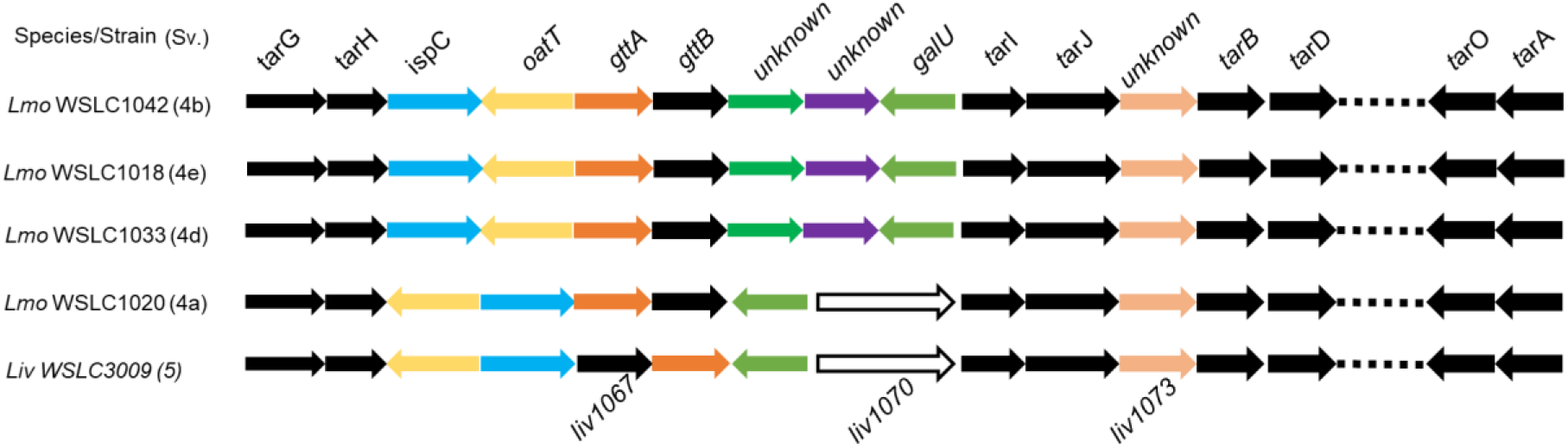
Comparison of genes encoding type II WTA biosynthesis pathways in the indicated *Listeria* serovars and strains.

**Table S1.** Homologs of Liv1070 proteins in a select number of *Listeria* subspecies (only top 20 are shown).

| Description | Scientific Name | Query Cover | E value | Per. ident | Acc. Len | Accession |
| --- | --- | --- | --- | --- | --- | --- |
| bifunctional glycosyltransferase family 2 protein/CDP-glycerol:glycerophosphate glycerophosphotransferase [*Listeria ivanovii*] | *Listeria ivanovii* | 100% | 0.00E+00 | 100 | 783 | WP_014092498.1 |
| bifunctional glycosyltransferase family 2 protein/CDP-glycerol:glycerophosphate glycerophosphotransferase [*Listeria ivanovii*] | *Listeria ivanovii* | 100% | 0.00E+00 | 99.87 | 783 | WP_111124699.1 |
| MULTISPECIES: bifunctional glycosyltransferase family 2 protein/CDP-glycerol:glycerophosphate glycerophosphotransferase [*Listeria*] | *Listeria* | 100% | 0.00E+00 | 97.06 | 783 | WP_003719308.1 |
| CDP-glycerol glycerophosphotransferase family protein [*Listeria ivanovii*] | *Listeria ivanovii* | 100% | 0.00E+00 | 96.81 | 783 | WP_185634543.1 |
| CDP-glycerol glycerophosphotransferase family protein [*Listeria seeligeri*] | *Listeria seeligeri* | 100% | 0.00E+00 | 96.55 | 783 | WP_194329958.1 |
| bifunctional glycosyltransferase family 2 protein/CDP-glycerol:glycerophosphate glycerophosphotransferase [*Listeria seeligeri*] | *Listeria seeligeri* | 100% | 0.00E+00 | 96.3 | 783 | WP_139590672.1 |
| CDP-glycerol glycerophosphotransferase family protein [*Listeria seeligeri*] | *Listeria seeligeri* | 100% | 0.00E+00 | 94.13 | 783 | WP_181249218.1 |
| CDP-glycerol glycerophosphotransferase family protein [*Listeria seeligeri*] | *Listeria seeligeri* | 100% | 0.00E+00 | 94 | 783 | WP_194341205.1 |
| CDP-glycerol glycerophosphotransferase family protein [*Listeria seeligeri*] | *Listeria seeligeri* | 100% | 0.00E+00 | 93.87 | 783 | WP_185365227.1 |
| CDP-glycerol glycerophosphotransferase family protein [*Listeria seeligeri*] | *Listeria seeligeri* | 100% | 0.00E+00 | 93.87 | 783 | WP_185338197.1 |
| CDP-glycerol glycerophosphotransferase family protein [*Listeria welshimeri*] | *Listeria welshimeri* | 100% | 0.00E+00 | 91.95 | 783 | WP_185303904.1 |
| unnamed protein product |  | 100% | 0.00E+00 | 91.95 | 783 | WP_194342659.1 |
| CDP-glycerol glycerophosphotransferase family protein [*Listeria welshimeri*] | *Listeria welshimeri* | 100% | 0.00E+00 | 91.95 | 783 | WP_194355605.1 |
| CDP-glycerol glycerophosphotransferase family protein [*Listeria welshimeri*] | *Listeria welshimeri* | 100% | 0.00E+00 | 91.95 | 783 | WP_194352431.1 |
| glycosyltransferase [*Listeria welshimeri*] | *Listeria welshimeri* | 100% | 0.00E+00 | 91.83 | 783 | MBC2346148.1 |
| glycosyltransferase [*Listeria welshimeri*] | *Listeria welshimeri* | 100% | 0.00E+00 | 91.83 | 783 | MBC1249545.1 |
| glycosyltransferase [*Listeria welshimeri*] | *Listeria welshimeri* | 100% | 0.00E+00 | 91.83 | 783 | MBC1611992.1 |
| glycosyltransferase [*Listeria welshimeri*] | *Listeria welshimeri* | 100% | 0.00E+00 | 91.83 | 783 | MBC1451016.1 |
| CDP-glycerol glycerophosphotransferase family protein [*Listeria welshimeri*] | *Listeria welshimeri* | 100% | 0.00E+00 | 91.83 | 783 | WP_194338958.1 |

**Table S2.**
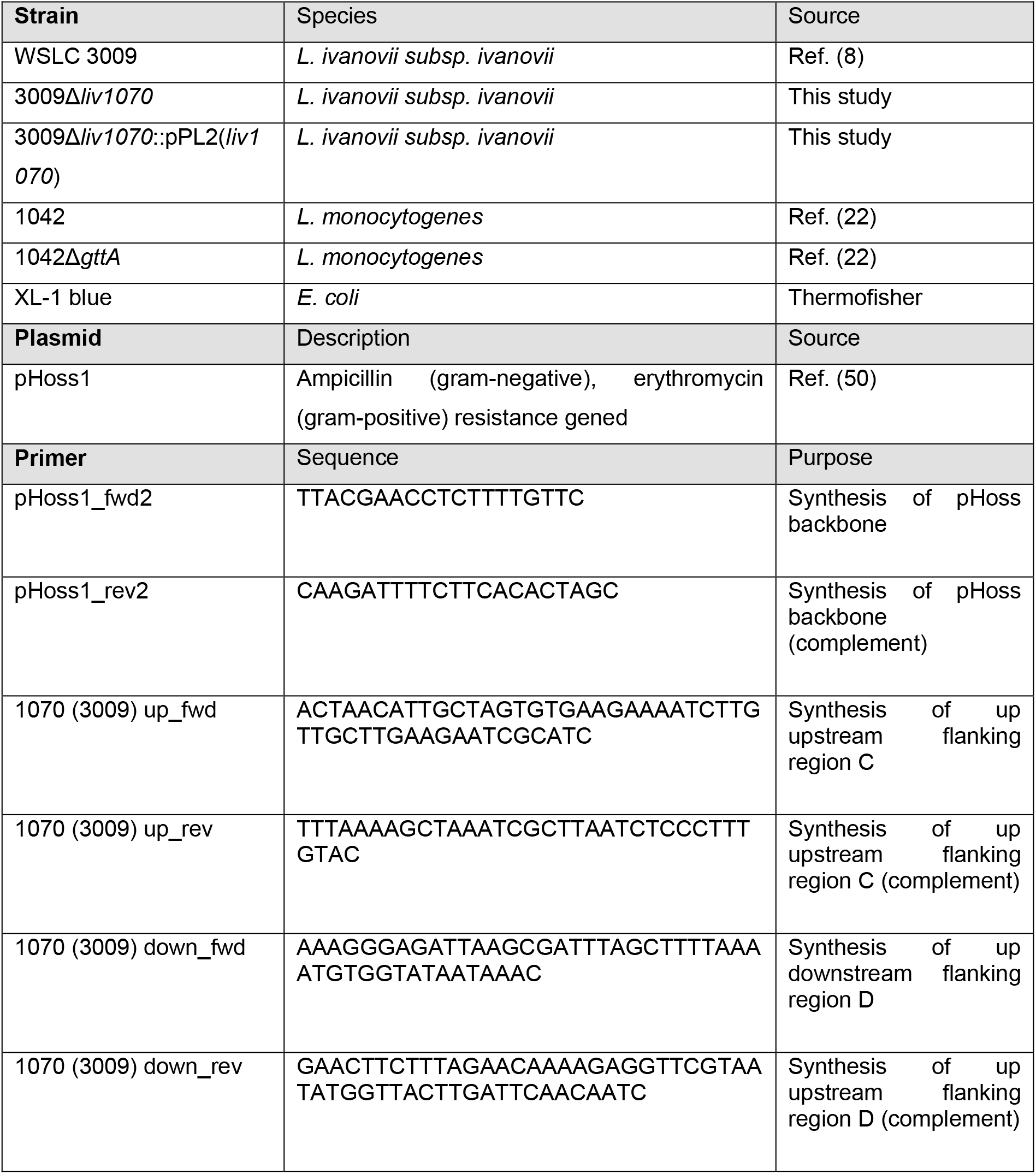
Srains, plasmids and primers.

## References

1. Charlier C, Perrodeau E, Leclercq A, Cazenave B, Pilmis B, Henry B, Lopes A, Maury MM, Moura A, Goffinet F, Dieye HB, Thouvenot P, Ungeheuer MN, Tourdjman M, Goulet V, de Valk H, Lortholary O, Ravaud P, Lecuit M, group Ms. 2017. Clinical features and prognostic factors of listeriosis: the MONALISA national prospective cohort study. Lancet Infect Dis 17:510–519.

2. Vazquez-Boland JA, Kuhn M, Berche P, Chakraborty T, Dominguez-Bernal G, Goebel W, Gonzalez-Zorn B, Wehland J, Kreft J. 2001. Listeria pathogenesis and molecular virulence determinants. Clin Microbiol Rev 14:584–640.

3. Gan L, Mao P, Jiang H, Zhang L, Liu D, Cao X, Wang Y, Wang Y, Sun H, Huang Y, Ye C. 2020. Two Prevalent Listeria ivanovii subsp. ivanovii Clonal Strains With Different Virulence Exist in Wild Rodents and Pikas of China. Front Vet Sci 7:88.

4. Elischerova K, Cupkova E, Urgeova E, Lysy J, Sesevickova A. 1990. [Isolation of Listeria ivanovii in Slovakia]. Cesk Epidemiol Mikrobiol Imunol 39:228–36.

5. Cummins AJ, Fielding AK, McLauchlin J. 1994. Listeria ivanovii infection in a patient with AIDS. J Infect 28:89–91.

6. Guillet C, Join-Lambert O, Le Monnier A, Leclercq A, Mechai F, Mamzer-Bruneel MF, Bielecka MK, Scortti M, Disson O, Berche P, Vazquez-Boland J, Lortholary O, Lecuit M. 2010. Human listeriosis caused by Listeria ivanovii. Emerg Infect Dis 16:136–8.

7. Snapir YM, Vaisbein E, Nassar F. 2006. Low virulence but potentially fatal outcome-Listeria ivanovii. Eur J Intern Med 17:286–7.

8. Hupfeld M, Trasanidou D, Ramazzini L, Klumpp J, Loessner MJ, Kilcher S. 2018. A functional type II-A CRISPR-Cas system from Listeria enables efficient genome editing of large non-integrating bacteriophage. Nucleic Acids Res 46:6920–6933.

9. Beye M, Gouriet F, Michelle C, Casalta JP, Habib G, Raoult D, Fournier PE. 2016. Genome analysis of Listeria ivanovii strain G770 that caused a deadly aortic prosthesis infection. New Microbes New Infect 10:87–92.

10. Orsi RH, den Bakker HC, Wiedmann M. 2011. Listeria monocytogenes lineages: Genomics, evolution, ecology, and phenotypic characteristics. Int J Med Microbiol 301:79–96.

11. Kamisango K, Fujii H, Okumura H, Saiki I, Araki Y, Yamamura Y, Azuma I. 1983. Structural and immunochemical studies of teichoic acid of Listeria monocytogenes. J Biochem 93:1401–9.

12. Fiedler F. 1988. Biochemistry of the cell surface of Listeria strains: a locating general view. Infection 16 Suppl 2:S92–7.

13. Brown S, Xia G, Luhachack LG, Campbell J, Meredith TC, Chen C, Winstel V, Gekeler C, Irazoqui JE, Peschel A, Walker S. 2012. Methicillin resistance in Staphylococcus aureus requires glycosylated wall teichoic acids. Proc Natl Acad Sci U S A 109:18909–14.

14. Winstel V, Kuhner P, Salomon F, Larsen J, Skov R, Hoffmann W, Peschel A, Weidenmaier C. 2015. Wall Teichoic Acid Glycosylation Governs Staphylococcus aureus Nasal Colonization. mBio 6:e00632.

15. Winstel V, Liang C, Sanchez-Carballo P, Steglich M, Munar M, Broker BM, Penades JR, Nubel U, Holst O, Dandekar T, Peschel A, Xia G. 2013. Wall teichoic acid structure governs horizontal gene transfer between major bacterial pathogens. Nat Commun 4:2345.

16. Wanner S, Schade J, Keinhorster D, Weller N, George SE, Kull L, Bauer J, Grau T, Winstel V, Stoy H, Kretschmer D, Kolata J, Wolz C, Broker BM, Weidenmaier C. 2017. Wall teichoic acids mediate increased virulence in Staphylococcus aureus. Nat Microbiol 2:16257.

17. Bielmann R, Habann M, Eugster MR, Lurz R, Calendar R, Klumpp J, Loessner MJ. 2015. Receptor binding proteins of Listeria monocytogenes bacteriophages A118 and P35 recognize serovar-specific teichoic acids. Virology 477:110–118.

18. Sumrall ET, Keller AP, Shen Y, Loessner MJ. 2020. Structure and function of Listeria teichoic acids and their implications. Mol Microbiol 113:627–637.

19. Uchikawa K, Sekikawa I, Azuma I. 1986. Structural studies on teichoic acids in cell walls of several serotypes of Listeria monocytogenes. J Biochem 99:315–27.

20. Shen Y, Boulos S, Sumrall E, Gerber B, Julian-Rodero A, Eugster MR, Fieseler L, Nystrom L, Ebert MO, Loessner MJ. 2017. Structural and functional diversity in Listeria cell wall teichoic acids. J Biol Chem 292:17832–17844.

21. Shen Y, Kalograiaki I, Prunotto A, Dunne M, Boulos S, Taylor NMI, Sumrall ET, Eugster MR, Martin R, Julian-Rodero A, Gerber B, Leiman PG, Menéndez M, Peraro MD, Cañada FJ, Loessner MJ. 2021. Structural basis for recognition of bacterial cell wall teichoic acid by pseudo-symmetric SH3b-like repeats of a viral peptidoglycan hydrolase. Chemical Science doi:10.1039/D0SC04394J.

22. Sumrall ET, Shen Y, Keller AP, Rismondo J, Pavlou M, Eugster MR, Boulos S, Disson O, Thouvenot P, Kilcher S, Wollscheid B, Cabanes D, Lecuit M, Grundling A, Loessner MJ. 2019. Phage resistance at the cost of virulence: Listeria monocytogenes serovar 4b requires galactosylated teichoic acids for InlB-mediated invasion. PLoS Pathog 15:e1008032.

23. Gessain G, Tsai YH, Travier L, Bonazzi M, Grayo S, Cossart P, Charlier C, Disson O, Lecuit M. 2015. PI3-kinase activation is critical for host barrier permissiveness to Listeria monocytogenes. J Exp Med 212:165–83.

24. Shen Y, Naujokas M, Park M, Ireton K. 2000. InIB-dependent internalization of Listeria is mediated by the Met receptor tyrosine kinase. Cell 103:501–10.

25. Sumrall ET, Schefer CRE, Rismondo J, Schneider SR, Boulos S, Grundling A, Loessner MJ, Shen Y. 2020. Galactosylated wall teichoic acid, but not lipoteichoic acid, retains InlB on the surface of serovar 4b Listeria monocytogenes. Mol Microbiol 113:638–649.

26. Carvalho F, Sousa S, Cabanes D. 2018. l-Rhamnosylation of wall teichoic acids promotes efficient surface association of Listeria monocytogenes virulence factors InlB and Ami through interaction with GW domains. Environ Microbiol 20:3941–3951.

27. Eugster MR, Morax LS, Huls VJ, Huwiler SG, Leclercq A, Lecuit M, Loessner MJ. 2015. Bacteriophage predation promotes serovar diversification in Listeria monocytogenes. Mol Microbiol 97:33–46.

28. Brown S, Santa Maria JP, Jr., Walker S. 2013. Wall teichoic acids of gram-positive bacteria. Annu Rev Microbiol 67:313–36.

29. Brown S, Meredith T, Swoboda J, Walker S. 2010. Staphylococcus aureus and Bacillus subtilis W23 make polyribitol wall teichoic acids using different enzymatic pathways. Chem Biol 17:1101–10.

30. Meredith TC, Swoboda JG, Walker S. 2008. Late-stage polyribitol phosphate wall teichoic acid biosynthesis in Staphylococcus aureus. J Bacteriol 190:3046–56.

31. Wang L, Lin M. 2008. A novel cell wall-anchored peptidoglycan hydrolase (autolysin), IspC, essential for Listeria monocytogenes virulence: genetic and proteomic analysis. Microbiology (Reading) 154:1900–1913.

32. Dorscht J, Klumpp J, Bielmann R, Schmelcher M, Born Y, Zimmer M, Calendar R, Loessner MJ. 2009. Comparative genome analysis of Listeria bacteriophages reveals extensive mosaicism, programmed translational frameshifting, and a novel prophage insertion site. J Bacteriol 191:7206–15.

33. Sumrall ET, Rohrig C, Hupfeld M, Selvakumar L, Du J, Dunne M, Schmelcher M, Shen Y, Loessner MJ. 2020. Glycotyping and specific separation of Listeria monocytogenes with a novel bacteriophage protein toolkit. Appl Environ Microbiol doi:10.1128/AEM.00612-20.

34. Vessey CJ, Wilding J, Folarin N, Hirano S, Takeichi M, Soutter P, Stamp GW, Pignatelli M. 1995. Altered expression and function of E-cadherin in cervical intraepithelial neoplasia and invasive squamous cell carcinoma. J Pathol 176:151–9.

35. Percy MG, Karinou E, Webb AJ, Grundling A. 2016. Identification of a Lipoteichoic Acid Glycosyltransferase Enzyme Reveals that GW-Domain-Containing Proteins Can Be Retained in the Cell Wall of Listeria monocytogenes in the Absence of Lipoteichoic Acid or Its Modifications. J Bacteriol 198:2029–42.

36. Banerjee M, Copp J, Vuga D, Marino M, Chapman T, van der Geer P, Ghosh P. 2004. GW domains of the Listeria monocytogenes invasion protein InlB are required for potentiation of Met activation. Mol Microbiol 52:257–71.

37. Khelef N, Lecuit M, Bierne H, Cossart P. 2006. Species specificity of the Listeria monocytogenes InlB protein. Cell Microbiol 8:457–70.

38. Lamond NM, Freitag NE. 2018. Vertical Transmission of Listeria monocytogenes: Probing the Balance between Protection from Pathogens and Fetal Tolerance. Pathogens 7.

39. Lamond NM, McMullen PD, Paramasvaran D, Visvahabrathy L, Eallanardo SJ, Maheswhari A, Freitag NE. 2020. Cardiotropic isolates of Listeria monocytogenes with enhanced vertical transmission dependent upon the bacterial surface protein InlB. Infect Immun doi:10.1128/IAI.00321-20.

40. Yin Y, Yao H, Doijad S, Kong S, Shen Y, Cai X, Tan W, Wang Y, Feng Y, Ling Z, Wang G, Hu Y, Lian K, Sun X, Liu Y, Wang C, Jiao K, Liu G, Song R, Chen X, Pan Z, Loessner MJ, Chakraborty T, Jiao X. 2019. A hybrid sub-lineage of Listeria monocytogenes comprising hypervirulent isolates. Nat Commun 10:4283.

41. Zhou M, Jiang M, Ren C, Liu S, Pu Q, Goldfine H, Shen H, Wang C. 2016. Listeria ivanovii Infection in Mice: Restricted to the Liver and Lung with Limited Replication in the Spleen. Front Microbiol 7:790.

42. Rismondo J, Haddad TFM, Shen Y, Loessner MJ, Grundling A. 2020. GtcA is required for LTA glycosylation in Listeria monocytogenes serovar 1/2a and Bacillus subtilis. Cell Surf 6:100038.

43. Ikeda M, Arai M, Okuno T, Shimizu T. 2003. TMPDB: a database of experimentally-characterized transmembrane topologies. Nucleic Acids Res 31:406–9.

44. Rismondo J, Percy MG, Grundling A. 2018. Discovery of genes required for lipoteichoic acid glycosylation predicts two distinct mechanisms for wall teichoic acid glycosylation. J Biol Chem 293:3293–3306.

45. Alvarez-Ordonez A, Leong D, Morgan CA, Hill C, Gahan CG, Jordan K. 2015. Occurrence, Persistence, and Virulence Potential of Listeria ivanovii in Foods and Food Processing Environments in the Republic of Ireland. Biomed Res Int 2015:350526.

46. Leong D, Alvarez-Ordonez A, Jordan K. 2014. Monitoring occurrence and persistence of Listeria monocytogenes in foods and food processing environments in the Republic of Ireland. Front Microbiol 5:436.

47. Oechslin F. 2018. Resistance Development to Bacteriophages Occurring during Bacteriophage Therapy. Viruses 10.

48. Shen Y, Loessner MJ. 2020. Beyond antibacterials - exploring bacteriophages as antivirulence agents. Curr Opin Biotechnol 68:166–173.

49. Lauer P, Chow MY, Loessner MJ, Portnoy DA, Calendar R. 2002. Construction, characterization, and use of two Listeria monocytogenes site-specific phage integration vectors. J Bacteriol 184:4177–86.

50. Abdelhamed H, Lawrence ML, Karsi A. 2015. A novel suicide plasmid for efficient gene mutation in Listeria monocytogenes. Plasmid 81:1–8.

